# The Genetic Chain Rule for Probabilistic Kinship Estimation

**DOI:** 10.1101/202879

**Authors:** Brian S. Helfer, Philip Fremont-Smith, Darrell O. Ricke

**Affiliations:** Bioengineering Systems and Technologies, MIT Lincoln Laboratory, Lexington, MA 02421 USA

**Keywords:** Forensic science, DNA forensics, single nucleotide polymorphism, SNP, kinship, kinship prediction

## Abstract

Accurate kinship predictions using DNA forensic samples has utility for investigative leads, remains identification, identifying relationships between individuals of interest, etc. High throughput sequencing (HTS) of STRs and single nucleotide polymorphisms (SNPs) is enabling the characterization of larger numbers of loci. Large panels of SNP loci have been proposed for improved mixture analysis of forensic samples. While multiple kinship prediction approaches have been established, we present an approach focusing on these large HTS SNP panels for predicting degree of kinship predictions. Formulas for first degree relatives can be multiplied (chained) together to model extended kinship relationships. Predictions are made using these formulations by calculating log likelihood ratios and selecting the maximum likelihood across the possible relationships. With a panel of 30,000 SNPs evaluated on an *in silico* dataset, this method can resolve parents from siblings and distinguish 1st, 2nd, and 3rd degree relatives from each other and unrelated individuals.

---

High throughput sequencing is revolutionizing capabilities in the fields of forensics, genetics, biology, and medicine. DNA forensics is evolving from sizing short tandem repeats (STRs) to sequencing STRs and single nucleotide polymorphisms (SNPs) (1, 2). Currently, DNA forensics uses STRs sized by capillary electrophoresis to perform individual identification, kinship predictions, and familial searching.

Familial searching is employed when the database being searched does not contain an exact match to a query STR profile. The use of identity by state (IBS), and/or likelihood ratio (LR) based searches enable query STR profiles to match potentially related individuals (3). In familial searching, likelihood ratio searches are often referred to as a kinship index (KI) (4). After relatives of the subject being searched are identified, lineage testing using mitochondrial DNA or Y chromosome STRs is typically performed (5) to confirm parental relationships. Familial searches are limited to first degree relatives due to the low number of STRs used (20 loci for US Combined DNA Index System - CODIS) (3), and the high probability of false positive matches when familial searching is expanded beyond first degree relations (6). Additional STR or SNP loci, and/or tools such as familias (7, 8), that use posterior probabilities to calculate likelihood ratios can be used to extend predictions beyond the first degree (9).

To increase kinship prediction capabilities, kinship prediction tools have been developed that use likelihood ratios, IBS, and SNP DNA microarrays (10, 11, 12, 13, 9). Several commercial companies like Ancestry.com, 23andMe, and Parabon leverage microarrays of SNPs for ancestry prediction with the capability of identifying close and distant relatives (14, 15). Huff et al. and Morimoto have used hundreds of thousands of SNPs characterized using SNP microarrays, to build conserved segments of chromosomal DNA, that are used to predict kinship (16, 17). Huff et al. reports 97% accuracy of to fifth degree relationships, but the selected parameters may not generalize across populations where the rate of linkage disequilibrium (LD) varies (18).

Although SNP microarrays can provide enhanced kinship prediction capabilities, they require 200ng of DNA (19), which is more than is typically available for forensic samples. As a result, interest in HTS technology, which can characterize SNPs with as little as 1 ng (20) of input DNA, has grown in the DNA forensics community. Most effort thus far has sought to use HTS SNP panels to improve resolution of complex mixtures (21, 22, 23). Data from these panels or new HTS kinship panels can be leveraged for kinship predictions. Mo *et al.* (24) selected an HTS panel of 472 SNPs for identifying second degree relationships using the likelihood ratio of autosomal matching. Machine learning and forensic HTS SNP mixture panels have been used to predict familial relationships across a set of three families in a tool named KinLinks (25). KinLinks trained a support vector machine based on features including the KING coefficient, identity by state (IBS), and identity by descent (IBD) (26). KinLinks was able to accurately predict all first degree and three quarters of second degree relationships. While machine learning models have the potential for accurate performance, they are highly dependent on the consistency of the training data, and are prone to over-optimization.

Here, we focus upon the kinship prediction between two individuals in the context of HTS panels designed for complex mixture analysis. The Genetic Chain Rule for Probabilistic Kinship Estimation provides a mathematical model that can predict likely relationship between two individuals (27). This work focuses on predicting degree of relatedness given a single profile against a large database of profiles for HTS SNP profiles. While KING coefficient and KI approaches provide insights into the closeness of relationships (28), this model attempts to predict the specific degree of relatedness between two individuals. This model does not require training data, thereby increasing generalizability. Furthermore, this work reflects the biological underpinnings of inheritance al-lowing for improved kinship predictions.

## Methods

### Input Data

All results were tested on an *in silico* dataset that simulated millions of individuals from African American, Estonian, Korean, and Palestinian ethnic groups. The data were simulated to provide an ideal representation of inheritance across multiple generations. The data were sampled to examine differences across varying degrees of relationship. A degree of relationship is defined by the expected percent of DNA shared between two individuals. For example, first degree relationships include parent-child, as well as siblings. In each case the individuals are expected to share approximately 50% of their DNA. Second degree relationships include grandparent-grandchild and aunt/uncle-niece/nephew where each share on average 25% of their DNA. Third degree relationships include cousins as well as great grandparent-great grandchild where approximately 12.5% of DNA is shared. The simulated data were produced using minor allele frequencies for 39,108 SNPs. These SNPs were selected as they were well characterized and included frequency information across the four selected groups as reported by the Allele Frequency Database (ALFRED) (29). A founding population of individuals were created for each ethnic group. The intermarriage rate was set to 0.15 for individuals with a single ethnicity and 1.0 for all other individuals. These individuals were assigned a gender and where randomly paired to create the next generation. The data were simulated across nine generations, with subsequent generations allowing for interbreeding between ethnic groups. This created varying levels of admixture with each new generation.

Non-admixed individuals from the last four generations of data and specific SNP groupings were used to assess individual relationships. Three SNP groups were subsets for each individual. SNP grouping were created by rank ordering SNPs with global minor allele frequencies (mAF) greater than or equal to 0.01 and selecting the top 2k, 20k, and 30k SNPs.

### Data Representation

All data are represented as a series of SNPs, with each locus having a minor allele, coupled with a mAF. The probability of the major allele occurring across a population is represented as *p*, while the probability of the minor allele occurring across a population is represented as *q*. As the SNPs analyzed have one major, and one minor allele, *p*, and *q* are set such that:

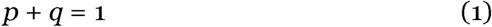

This follows Hardy-Weinberg equilibrium allowing for the frequency of genotypes to be represented as *p*^2^, 2*pq*, or *q*^2^. By satisfying Eq. 1, it is ensured that:

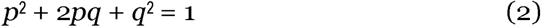

Using this information, the conditional probability of any genotype occurring, given another individual of a known relationship having a certain genotype can be derived. The genotype of an individual with parents from different populations (designed with subscripts 1 and 2) and admixed individuals are modeled with:

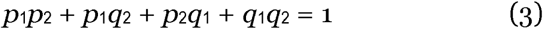

### Child Given Known Parent

The first relationship calculated is the probability of a child having a particular genotype *G*_*c*_, given that the parent has a genotype *G*_*p*1_. Given that one parent’s genotype is known, the possible alleles that they could pass on to their child is also known. The probability of inheriting each allele from *G*_*p*1_ for the child is 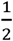. The genotype of the second parent is represented as *G*_*p*2_.

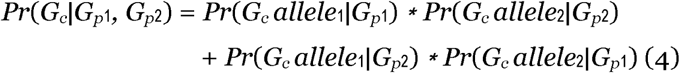

The highest rate of de novo human germline mutations occurs at a rate of 1.3×10^*-*7^ for methylated CpG dinucleotides (30); this rate is too low to affect most individual HTS SNP profiles. A very small number of impossible parent to child SNP loci are tolerated to accommodate possible HTS sequencing errors.

### Parent Given Known Child

Leveraging the information presented in Table 1, it is possible to calculate the probability of a parent having a particular genotype *G*_*p*_, given that the child has a known genotype *G*_*c*_. This is formulated through an application of Bayes’ rule, and is presented in Table 2.

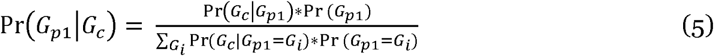

**Table 1.** Probability of the event (*α*) an individual with a given genotype (*Ind*_*α*_), conditioned (*β*) on another individual (*Ind*_*β*_) having a given genotype. The genotype letter (**A**) represents the major allele with population frequency *p*_*i*_ for individual i, while genotype letter (**a**) represents the minor allele with population frequency *q*_*i*_; this allows individuals to have different ethnicities.

| $Ind_\alpha$ | $Ind_\beta$ | $\Pr(Ind_\beta = Child \vert Ind_\alpha = Parent_1)$<br>Parent<sub>1</sub> : $p_1, q_1$,<br>Parent<sub>2</sub> : $p_2, q_2$ | $\Pr(Ind_\beta = Parent_1 \vert Ind_\alpha = Child)$<br>Parent<sub>1</sub> : $p_1, q_1$,<br>Parent<sub>2</sub> : $p_2, q_2$ |
| --- | --- | --- | --- |
| AA | AA | $p_2$ | $p_1$ |
| AA | Aa | $q_2$ | $q_1$ |
| AA | aa | 0 | 0 |
| Aa | AA | $0.5p_2$ | $\frac{q_2 p_1^2}{p_1 q_2 + q_1 p_2}$ |
| Aa | Aa | $0.5p_2 + 0.5q_2 = 0.5$ | $\frac{p_1 q_1}{p_1 q_2 + q_1 p_2}$ |
| Aa | aa | $0.5q_2$ | $\frac{p_2 q_1^2}{p_1 q_2 + q_1 p_2}$ |
| aa | AA | 0 | 0 |
| aa | Aa | $p_2$ | $p_1$ |
| aa | aa | $q_2$ | $q_1$ |
| $Ind_\alpha$ | $Ind_\beta$ | $\Pr(Ind_\beta = child_1 \vert Ind_\alpha = child_2)$<br>Parent<sub>1</sub> : $p_1, q_1$,<br>Parent<sub>2</sub> : $p_2, q_2$ | $\Pr(Ind_\beta = Grandchild \vert Ind_\alpha = Grandparent_1)$<br>Parent<sub>1</sub> : $p_1, q_1$,<br>Parent<sub>2</sub> : $p_2, q_2$,<br>Grandparent<sub>1</sub> : $p_3, q_3$,<br>Grandparent<sub>2</sub> : $p_4, q_4$ |
| AA | AA | $p_1 p_2 + 0.5 p_1 q_2 + 0.5 q_1 p_2 + 0.25 q_1 q_2$ | $p_2 p_4 + 0.5 p_2 q_4$ |
| AA | Aa | $0.5(p_1 q_2 + q_1 p_2 + q_1 q_2)$ | $q_2 p_4 + 0.5 q_4$ |
| AA | aa | $0.25 q_1 q_2$ | $0.5 q_2 q_4$ |
| Aa | AA | $0.5(p_1 p_2 + \frac{p_1 q_1 p_2 q_2}{p_1 q_2 + q_1 p_2})$ | $0.5 p_2 p_4 + 0.25 p_2$ |
| Aa | Aa | $0.5(p_1 p_2 + q_1 q_2) + \frac{p_1^2 q_2^2 + p_1 q_1 p_2 q_2 + q_1^2 p_2^2}{p_1 q_2 + q_1 p_2}$ | $0.5 p_2 q_4 + 0.25 + 0.5 q_2 p_4$ |
| Aa | aa | $0.5(q_1 q_2 + \frac{p_1 q_1 p_2 q_2}{p_1 q_2 + q_1 p_2})$ | $0.25 q_2 + 0.5 q_2 q_4$ |
| aa | AA | $0.25 p_1 p_2$ | $0.5 p_2 p_4$ |
| aa | Aa | $0.5(p_1 q_2 + q_1 p_2 + p_1 p_2)$ | $0.5 p_4 + p_2 q_4$ |
| aa | aa | $q_1 q_2 + 0.5 p_1 q_2 + 0.5 q_1 p_2 + 0.25 p_1 p_2$ | $0.5 p_2 q_4 + q_2 q_4$ |

**Table 2.**
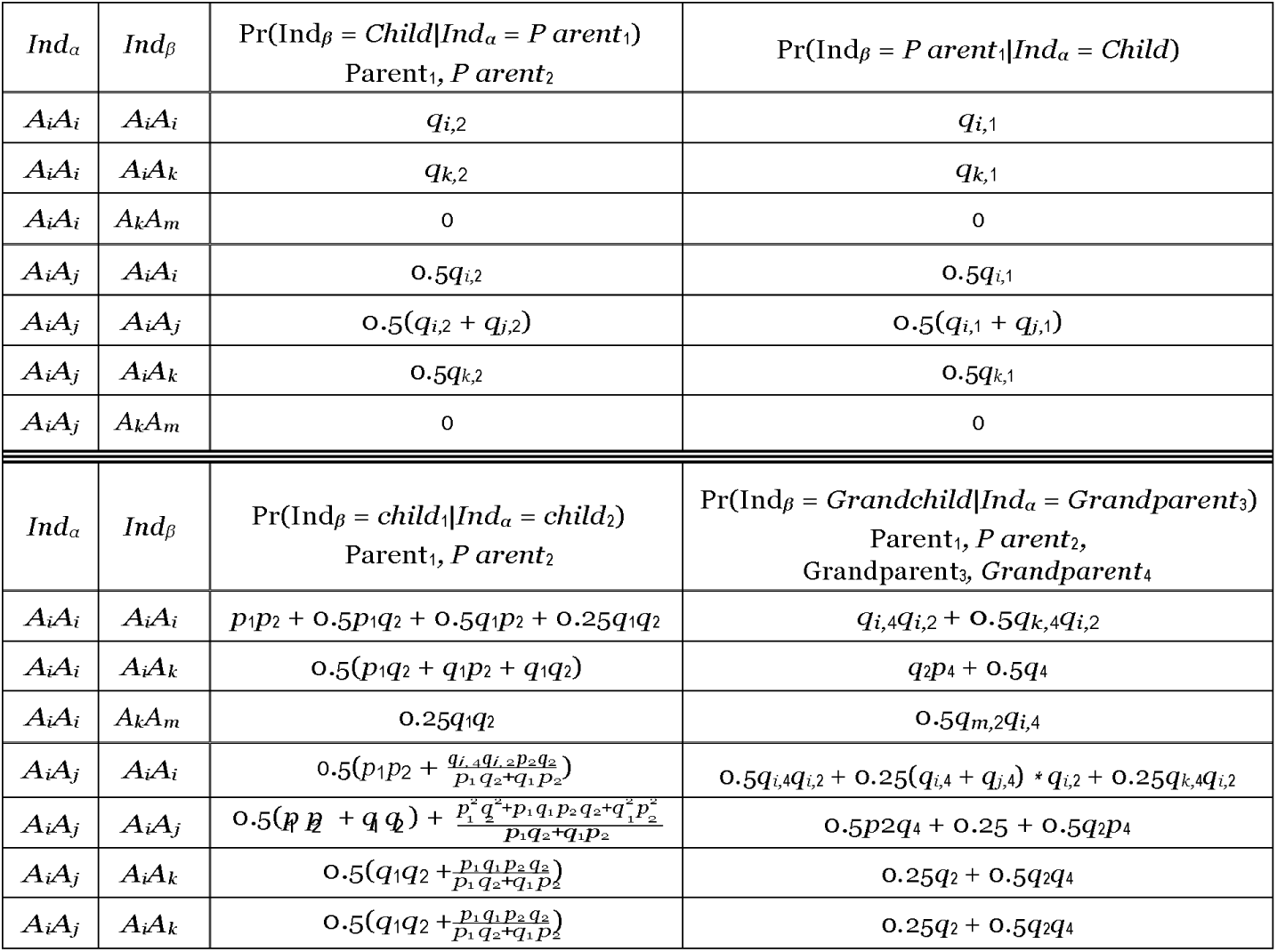
Probability of the event (*α*) an individual with a given genotype (*Ind*_*α*_), conditioned (*β*) on another individual (*Ind*_*β*_) having a given genotype. The genotype letter (**A**) represents the major allele with population frequency *p*_*i*_ for individual i, while genotype letter (**a**) represents the minor allele with population frequency *q*_*i*_; this allows individuals to have different ethnicities.

In the above and subsequent equations, *G*_*i*_, is used to represent all possible allele combinations. As a result, *G*_*i*_ can be expressed as:

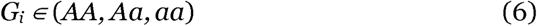

When the both parents and children have the same ethnic background, it is possible to show that *Pr*(*G*_*p*1_|*G*_*c*_) = *Pr*(*G*_*c*_|*G*_*p*1_)

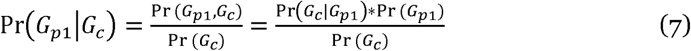

As *P r*(*G*_*c*_) is equal to *P r*(*G*_*p*1_) it is shown that *P r*(*G*_*p*1_|*G*_*c*_) = *P r*(*G*_*c*_|*G*_*p*1_).

### Sibling Relationships

It is possible to use this information to further compute the probability that a child will have a genotype *G*_*c*1_ given that a sibling of theirs has an observed genotype *G*_*c*2_. In order to properly compute the probability of genotype *G*_*c*1_ occurring, it is essential to factor in genotypes of the two parents *G*_*p*1_, and *G*_*p*2_. Using this information, the desired sibling-sibling conditional probability is computed as the probability of Child 1 having a genotype *G*_*c*1_ given the possible genotypes that their parents could have, multiplied by the probability of the two parents having genotypes *G*_*p*1_ and *G*_*p*2_ given the known genotype of Child 2.

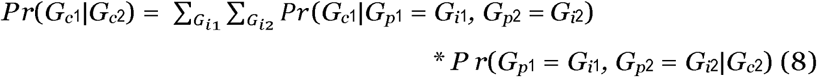

Given an assumption that the two parents are not close relatives, it is possible to further rewrite Eq. 6 as:

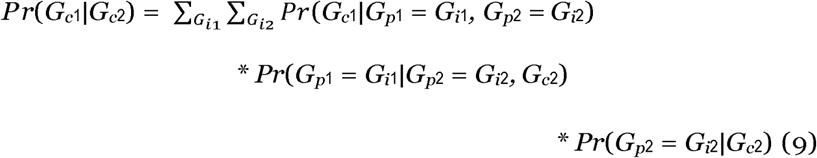

As shown in Eq. 7, it becomes possible to compute the probability of a child having a particular genotype given that their sibling has a known genotype. This is done by computing the product of the probability of a child having genotype *G*_*c*1_ normalized across the genotypes for both parents *G*_*p*1_, and *G*_*p*2_, and the probability of each parent having a genotype given the knowledge of the second child’s genotype. This equation is the product of three terms, where the first term represents the probability of a child having a genotype given specified genotypes for their parents, the second term represents the probability of the first parent having a specified genotype given the child’s sibling and their other parent’s genotype, and the third term represents the probability of the second parent having a particular genotype given the child’s sibling.

### Bayesian Chain (Product) Rule of Kinship

The above framework can be generalized to compute the probability of a particular genotype given any relationship between two people. This formulation is defined as the Bayesian Chain (Product) Rule of Kinship. The Bayesian Chain Rule of Kinship expresses any relationship between individuals as the **product** of a series of relationships. For instance, if one wished to compute the cousin relationship between *M*_31_ and *F*_33_ as shown in Figure 1, one would represent this as the relationship between child and parent, parent and sibling, and the parent’s sibling and their child. As can be noted, all components of the chain rule take the form of child given parent to move up the tree, sibling given sibling to move across the tree, and parent given child to move down the tree. These operations allow for complete navigation between any two individuals. Expressed another way:

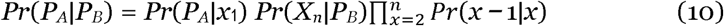

**FIG. 1.**
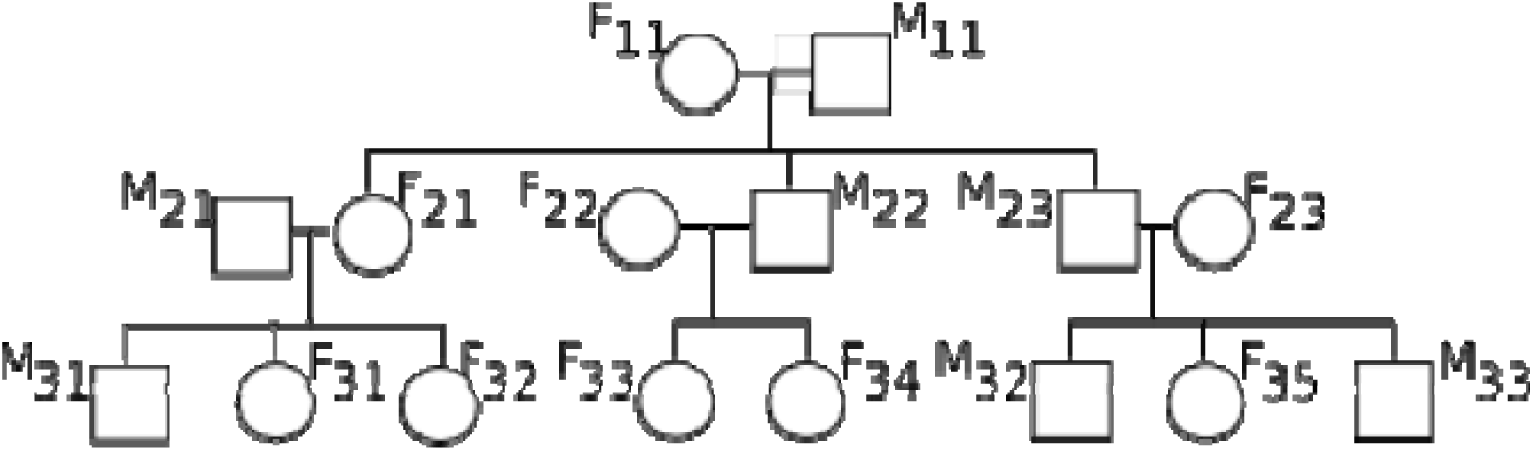
Reference Family Tree. Each individual is represented as a female (F) or male (M) along with a two digit number representing their generation, followed by a unique identifier for that person within the generation.

This takes the product of all people between two individuals, and uses the Bayesian Chain Rule of Kinship to compute a probability of a genotype given a particular relationship.

### Extended Relationships

Extended relationships can be computed using the previously defined Bayesian Chain Rule of Kinship. Figure 1 shows a family tree where each individual is identified as male (M) or female (F), and with two indices identifying their generation, along with a unique identifier for that individual within the generation. For instance, *F*_23_ represents the third unique woman appearing in the second generation.

### Grandchild Given Grandparent

The probability of a child (*M*_31_) having a given genotype *G*_*c*_, given their grandparent (*M*_11_) has a known genotype *G*_*g*_ can be computed using the Markov and chain rule assumptions to model the child (*M*_31_) as dependent on their parent (*F*_21_), and the parent (*F*_21_) to be dependent on the grandparent (*M*_11_).

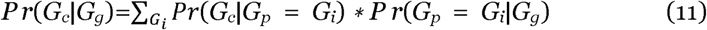

Since child (*M*_31_) inherits DNA from parent (*F*_21_), and parent (*F*_21_) inherits DNA from grandparent (*M*_11_), the probability equation decomposes into the child being directly dependent on their parent, and the parent being directly de-pendent on the grandparent. It is unnecessary to condition the child’s genotype on the grandparent’s genotype, as that is already factored into the parent’s genotype. Given that other parent has an unknown genotype, *G*_*i*_ is used to marginalize over all possible genotypes for that parent.

### Child Given Aunt/Uncle

The same principles apply to identify the likelihood that a child will have a genotype given that their aunt/uncle have a known genotype *G*_*au*_. In this case, the probability of the child’s genotype is decomposed into the relationship between child and parent, and parent and sibling.

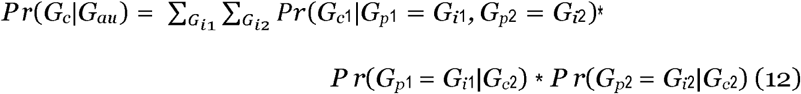

### Log Likelihood Calculation

The above formulations can be further used to calculate the log likelihood of two individuals having a particular relationship given the observed data. The log likelihood is defined as the probability of data (D) given a hypothesis (H).

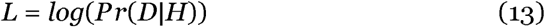

In familial identification, the combined probability of observing a set of SNPs given an assumed relationship is computed by taking the product of all conditional probabilities across SNPs. However, for large SNP panels this value can become difficult to store in memory without seeing significant truncation of the likelihood. For this reason, the sum of *log* of the probabilities is calculated at each SNP position. The log likelihood is computed in this manner for each of the test relationships. The predicted relationship is then given as the *argmax* between all the expected relationships and an unrelated individual. This allows for the computation of the likelihood of any relationship given the observed genotypes of two individuals.

### Linkage Disequilibrium

In order to account for SNP linkage disequilibrium, an adjustment factor for L based on the linkage disequilibrium D’ value between two linked SNP loci is applied. If the SNPs are unlinked D’ is zero with no adjustment of L. If the SNPs are highly linked, L is adjusted by (1-D’). Adjacent SNPs with full linkage disequilibrium (e.g., D’=1) resulted in prediction scoring for the first SNP only. Adjacent SNPs with partial linkage disequilibrium had prediction scores for the next SNP adjusted by (1-D’).

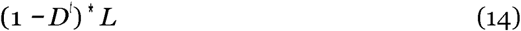

### Current Limitations

The current calculations rely on the independence of inheritance of all alleles. This simplifies the calculation; however, it does not account for haploblocks, or sex chromosomes. As a result, it is not possible to distinguish between different relationships that are two or more generations apart, or the direction of the relationship. This study presents a panel of 30k SNPs of assumed independence that is useful for the theoretical modeling of panel size contribution, and for organisms that have a larger genome and this number of independent SNPs is possible. This work provides a framework that is generalizable and can be extended in the future to incorporate haploblocks and sex chromosomes.

## Results

The previously defined mathematical relationships were validated using the previously described *in silico* database of four ethnicities and the lower four of the nine generations. The data was further subdivided into four different ethnic groups which have separate mAF values across the 39,108 sampled SNPs.

### Data Relationship Separability

The relationship separability was examined as a function of the number of differences across SNPs. A difference was defined as the number of discordant alleles at each locus with a value between zero and two. The number of discrepancies was summed across all SNPs for a single pairwise relationship. This was then done for one thousand examples of each relationship. A kernel density estimate was fitted to this distribution and then shown in the figures below. The kernel density estimate was computed using the Python package seaborn, which fits a series of distributions across the set of points and then smooths them to make a single distribution.

The number of differences across degree were plotted while varying the number of SNPs used in the comparison. Figure 2 plots differences across distributions using the full panel of 30k SNP loci. The level of separation is then plotted for 20k SNPs as shown in Figure 3, and finally the number of differences is examined with the panel reduced to only 2k SNPs, as shown in Figure 4.

**FIG. 2.**
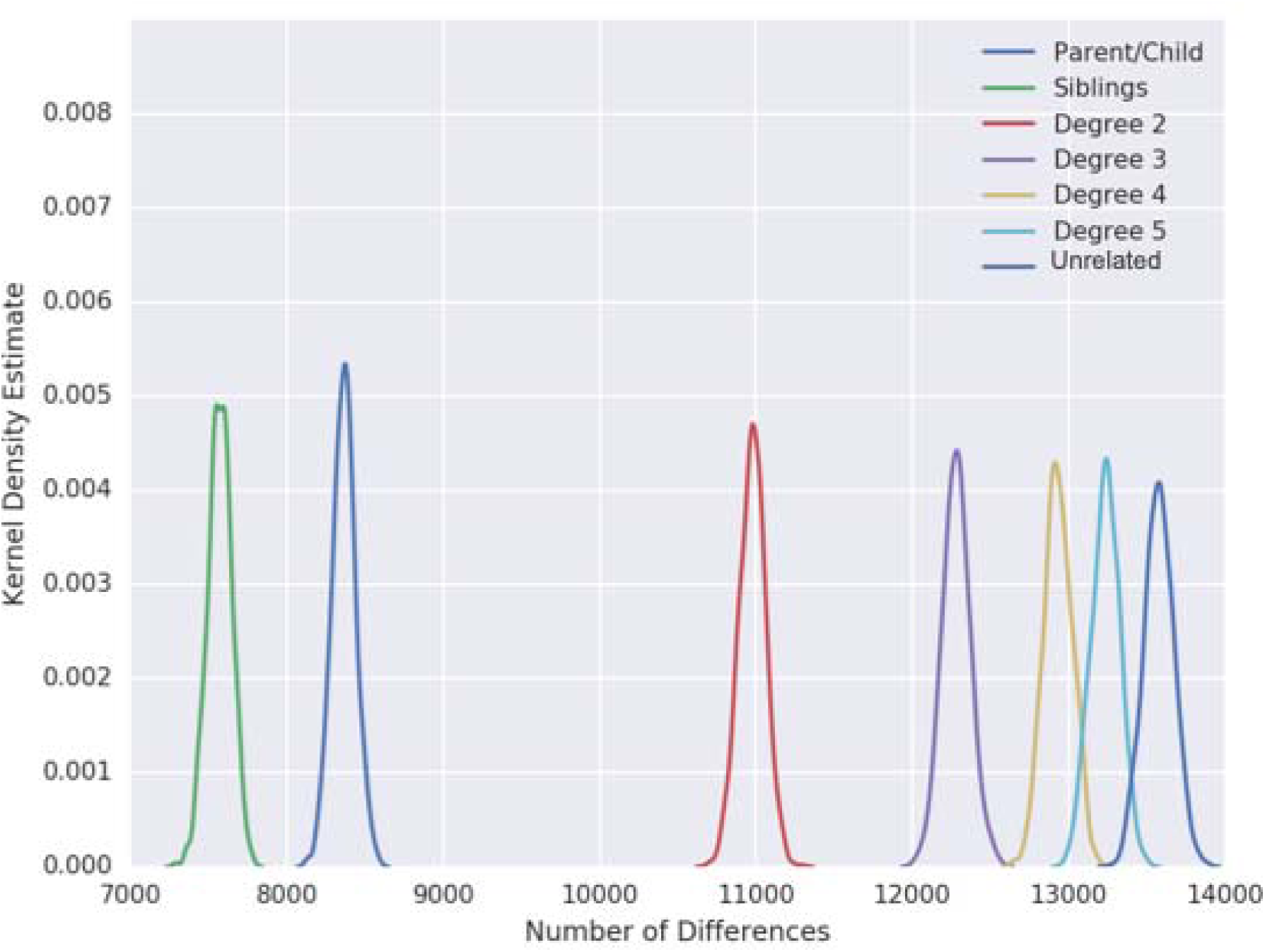
Differences separated by degree of relationship for a panel of 30k SNPs. The number of differences across pairwise relationships are plotted as a kernel density estimate. Peaks are separated up to and including 3rd degree, while 4th, and unrelated individuals show minimal overlap.

**FIG. 3.**
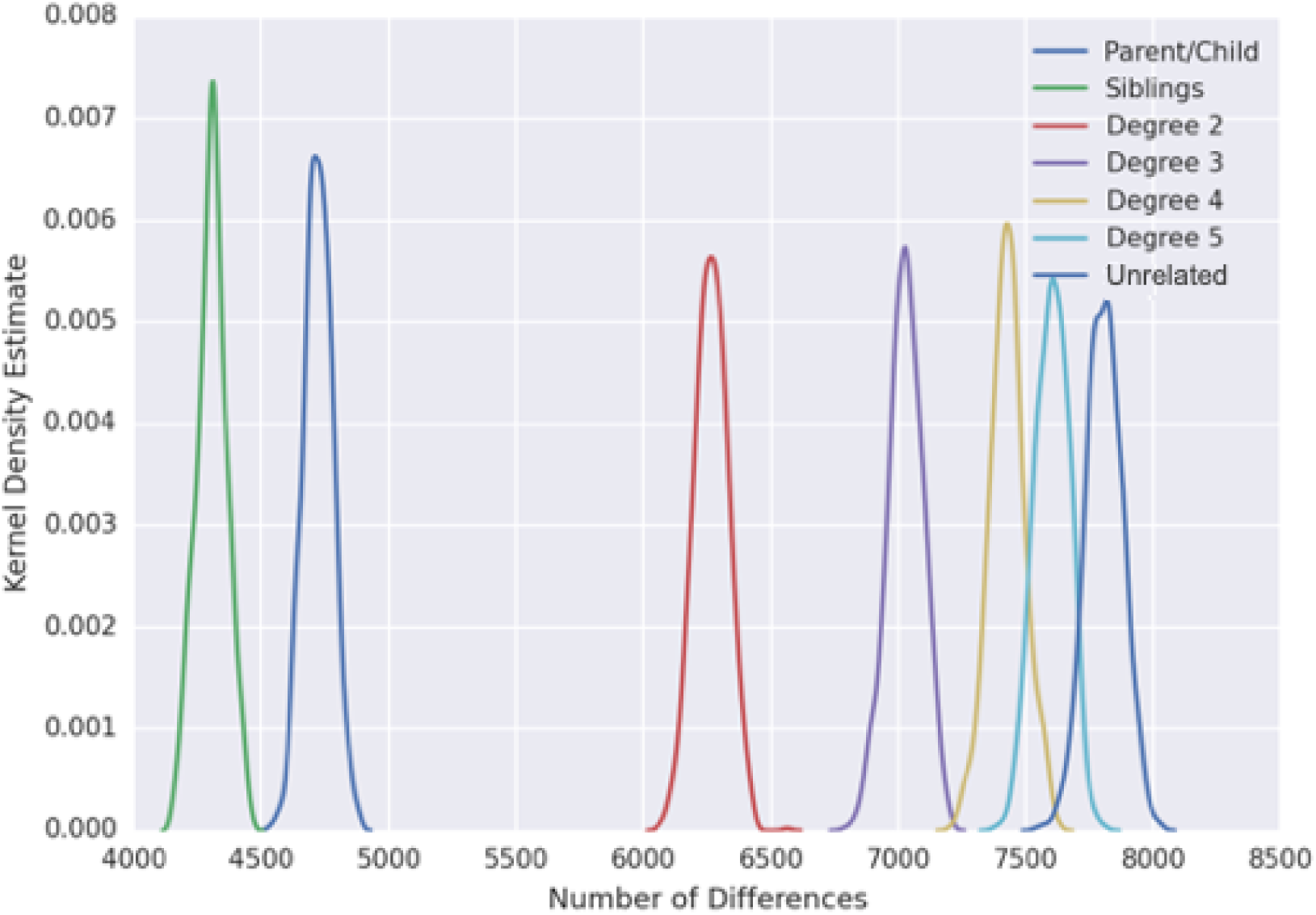
Differences separated by degree of relationship for a panel of 20k SNPs. The number of differences across pairwise relationships are plotted as a kernel density estimate.

**FIG. 4.**
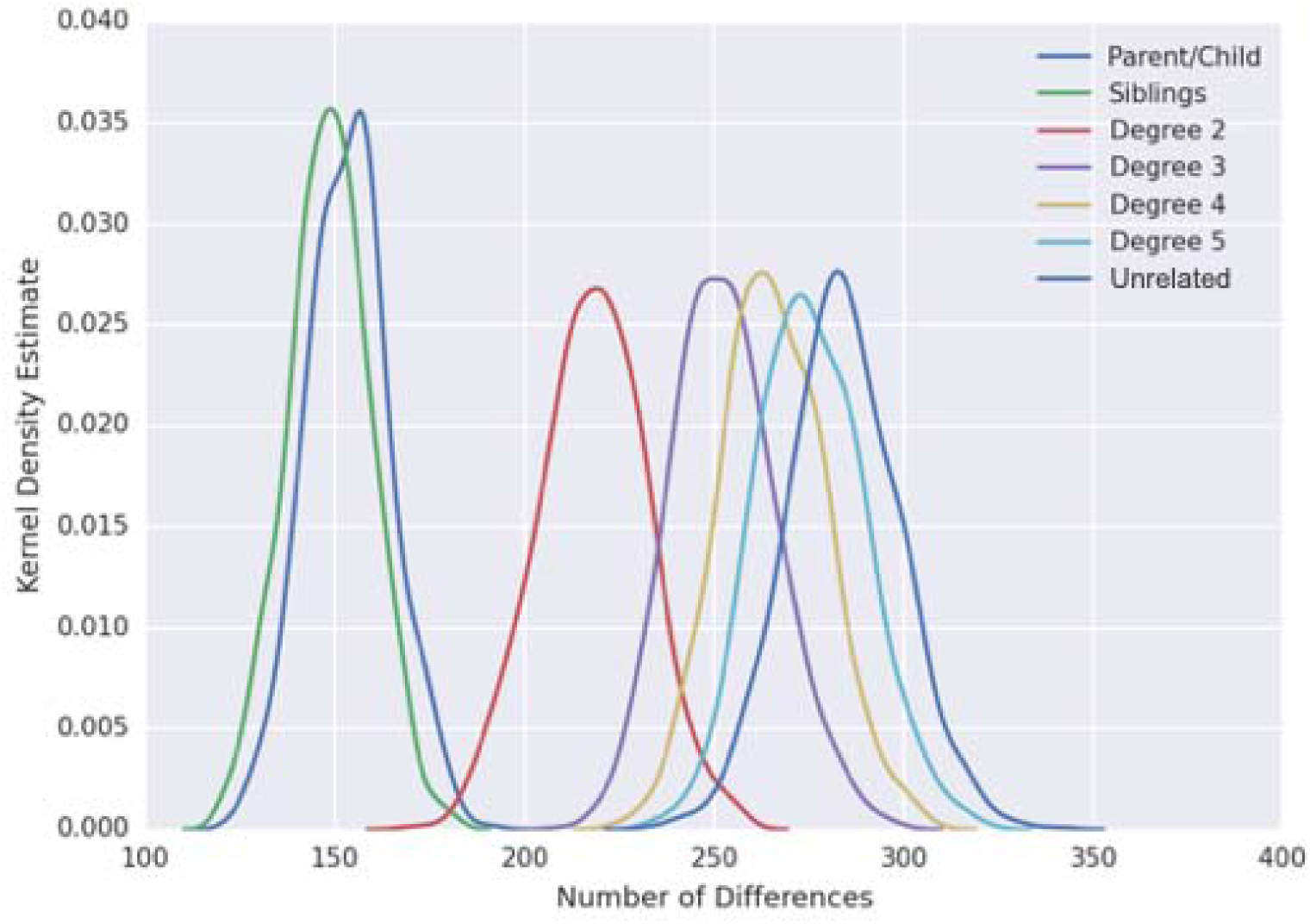
Differences separated by degree of relationship for a panel of 2k SNPs. The number of differences across pairwise relationships are plotted as a kernel density estimate.

### Log Likelihood Prediction

After examining differences between individuals, the log likelihood was then used to predict the degree of relation across pairs of individuals. At each pair, the algorithm identifies if it is a parent-child relationship, a sibling-sibling relationship, 2^*nd*^ to 5^*th*^ degree relationship, or two unrelated individuals. The performance for this assessment is shown in Table 3. These results present a theoretical optimum for performance given the current algorithm. The next step is the application of this approach to families with extended family members. These results are useful as templates for differences in performance for SNP panels with different number of loci.

**Table 3.** Confusion matrix for 30k SNP panel degree prediction. Rows of the matrix show the true number of relationships, while the columns represent the predicted number of relationships across pairings.

|  | Parent-Child | Sibling | Degree 2 | Degree 3 | Degree 4 | Degree 5 | Unrelated |
| --- | --- | --- | --- | --- | --- | --- | --- |
| Parent-Child | 2000 | 0 | 0 | 0 | 0 | 0 | 0 |
| Sibling | 0 | 1000 | 0 | 0 | 0 | 0 | 0 |
| Degree 2 | 0 | 0 | 4000 | 0 | 0 | 0 | 0 |
| Degree 3 | 0 | 0 | 0 | 5000 | 0 | 0 | 0 |
| Degree 4 | 0 | 0 | 0 | 1 | 995 | 5 | 0 |
| Degree 5 | 0 | 0 | 0 | 0 | 5 | 993 | 2 |
| Unrelated | 0 | 0 | 0 | 0 | 0 | 2 | 998 |

## Discussion

In this study, *in silico* data were used to identify the degree of relatedness between individuals spanning four generations. HTS sequencing of large SNP panels has been demonstrated by for identification and mixture analysis (21, 22, 31). MIT Lincoln Laboratory has tested SNP panels up to 3,500 (31) and 15,000 SNPs with current AmpliSeq panel designs of up to 24,000 SNPs possible. The 2k and 30k SNP panels used in this study selected SNPs with lower mAF for consistency with SNP panels optimized for mixture analysis. As the number of SNPs increase above 20k, linkage disequilibrium between SNPs will be encountered. Figures 2 and 3 demonstrate a clear separability between parent-child relationships and siblings, as well as between individuals with second, third, and unrelated levels of relationship. This was reflected by the use of log likelihood values to fully and correctly identify the difference between individuals of these different degrees. As the degree increases, the curves become closer together. The upper tail of the 4th degree relatives is near the lower tail of unrelated individuals. For the 20k and 30k SNP panels, the distribution for 5th degree relatives overlap the distributions for 4th degree relatives and unrelated individuals. The confusion matrix shown in Table 3 for this method illustrates the high accuracy on these *in silico* pedigrees. The impact of reducing the SNP panel size is illustrated in Figure 3 for 20k SNPs and Figure 4 for 2k SNPs. For the 2k SNPs panel, the different relationships become much less separable with substantial overlap seen at 3rd, 4th, 5th, and unrelated individuals. These results are comparable to other estimates of kinship along numerical scales like KI, *etc*. (32, 33, 28) with the advantage of predicting the specific degree of kinship for two individuals with application for low DNA HTS forensic samples.

This method is complimentary to other techniques that build predictors (25) based on identity by descent, or likelihood models based off of conserved chromosomal segments (16). The genetic chain rule presented in this paper is capable of sampling at separated loci with minimal or very low linkage disequilibrium (LD). Beyond a SNP spacing of 160,000 base pairs, the LD *r*^2^ (or D’) values are typically less than 0.1 (18).

## Conclusion

In this work, we present a Bayesian framework for identifying the level of relation between different individuals. This framework builds on the biology of inheritance, along with statistical calculation to predict degree of relation without requiring a training database or parameter optimization. This allows for further improvement by incorporating more biological properties into the model.

## Acknowledgements

The authors would like to thank Anthony Trasatti and James Watkins for useful discussions.

This material is based upon work supported under Air Force Contract No. FA8702-15-D-0001. Any opinions, findings, conclusions, or recommendations expressed in this material are those of the author(s) and do not necessarily reflect the views of the U.S. Air Force.

## References

1. B. Sobrino, M. Brión, A. Carracedo, SNPs in forensic genetics: a review on SNP typing methodologies, Forensic Sci Int 154 (2) (2005) 181–194.

2. A. J. Pakstis, W. C. Speed, J. R. Kidd, K. K. Kidd, Candidate SNPs for a universal individual identification panel, Human Genet 121 (3-4) (2007) 305.

3. J. Ge, R. Chakraborty, A. Eisenberg, B. Budowle, Comparisons of familial DNA database searching strategies, J Forensic Sci 56 (6) (2011) 1448–1456.

4. J. Ge, B. Budowle, Kinship index variations among populations and thresholds for familial searching, PLOS ONE 7 (5) (2012) 1–8.

5. H. Liu, X. Li, J. Mulero, A. Carbonaro, M. Short, J. Ge, A convenient guideline to determine if two Y-STR profiles are from the same lineage, Electrophoresis 37 (12) (2016) 1659–1668.

6. E. Niedzwiecki, S. Debus-Sherrill, M. B. Field, Understanding Familial DNA Searching: Coming to a Consensus on Terminology, ICF Inter, 2016.

7. D. Kling, A. O. Tillmar, T. Egeland, Familias 3 – extensions and new functionality, Forensic Science International: Genetics 13 (2014) 121–127.

8. T. Egeland, P. Mostad, B. Mevåg, M. Stenersen, Beyond traditional paternity and identification cases: Selecting the most probable pedigree, Forensic Sci Inter 110 (1) (2000) 47–59.

9. D. Kling, T. Egeland, M. H. Piñero, M. D. Vigeland, Evaluating the statistical power of DNA-based identification, exemplified by ‘the missing grandchildren of argentina’, Forensic Sci Int Genet 31 (2017) 57–66.

10. D. Kling, J. Welander, A. Tillmar, Øivind Skare, T. Egeland, G. Holmlund, DNA microarray as a tool in establishing genetic relatedness—current status and future prospects, Forensic Sci Inter Genet 6 (3) (2012) 322–329.

11. D. Kling, SNP analyzer - a tool to analyze large sets of genetic markers accounting for linkage, Forensic Sci Int Genet Suppl 6 (2017) e587–e588.

12. V. Heinrich, T. Kamphans, S. Mundlos, P. N. Robinson, P. M. Krawitz, A likelihood ratio-based method to predict exact pedigrees for complex families from next-generation sequencing data, Bioinformatics 33 (1) (2016) 72–78.

13. J. Huisman, Pedigree reconstruction from SNP data: parentage assignment, sibship clustering and beyond, Mol Ecology Resources 17 (5) (2017) 1009–1024.

14. P. J. Zettler, J. S. Sherkow, H. T. Greely, 23andMe, the food and drug administration, and the future of genetic testing, JAMA Internal Med 174 (4) (2014) 493–494.

15. H. Wolinsky, CSI on steroids: DNA-based phenotyping is helping police derive visual information from crime scene samples to aid in the hunt for suspects, EMBO Reports 16 (7) (2015) 782–786.

16. C. D. Huff, D. J. Witherspoon, T. S. Simonson, J. Xing, W. S. Watkins, Y. Zhang, et al., Maximum-likelihood estimation of recent shared ancestry (ERSA), Genome Res 21 (5) (2011) 768–774.

17. C. Morimoto, S. Manabe, S. Fujimoto, Y. Hamano, K. Tamaki, Discrimination of relationships with the same degree of kinship using chromosomal sharing patterns estimated from high-density SNPs, Forensic Sci Int Genet 33 (2018) 10–16.

18. S. Shifman, J. Kuypers, M. Kokoris, B. Yakir, A. Darvasi, Linkage disequilibrium patterns of the human genome across populations, Human Mol Genet 12 (7) (2003) 771–776.

19. M. A. Quail, M. Smith, P. Coupland, T. D. Otto, S. R. Harris, T. R. Connor, et al., A tale of three next generation sequencing platforms: comparison of Ion Torrent, Pacific Biosciences and Illumina MiSeq sequencers, BMC Genomics 13 (1) (2012) 341.

20. A. Adey, H. G. Morrison, X. Xun, J. O. Kitzman, E. H. Turner, B. Stackhouse, et al., Rapid, low-input, low-bias construction of shotgun fragment libraries by high-density in vitro transposition, Genome Biol 11 (12) (2010) R119.

21. L. Voskoboinik, A. Darvasi, Forensic identification of an individual in complex DNA mixtures, Forensic Sci Int Genet 5 (2010) 428–435.

22. J. Isaacson, E. Schwoebel, A. Shcherbina, D. Ricke, J. Harper, M. Petrovick, et al., Robust detection of individual forensic profiles in DNA mixtures, Forensic Sci Int Genet 14 (2015) 31–37.

23. H.-L. Hwa, W.-C. Chung, P.-L. Chen, C.-P. Lin, H.-Y. Li, H.-I. Yin, et al., A 1204-single nucleotide polymorphism and insertion-deletion polymorphism panel for massively parallel sequencing analysis of DNA mixtures, Forensic Sci Int Genet 32 (2018) 94–101.

24. S.-K. Mo, Z.-L. Ren, Y.-R. Yang, Y.-C. Liu, J.-J. Zhang, H.-J. Wu, et al., A 472-SNP panel for pairwise kinship testing of second-degree relatives, Forensic Sci Int Genet 34 (2018) 178–185.

25. A. Shcherbina, D. O. Ricke, E. Schwoebel, T. Boettcher, C. Zook, J. Bobrow, et al., KinLinks: Software toolkit for kinship analysis and pedigree generation from HTS datasets, in: 2016 IEEE International Symposium on Technologies for Homeland Security (HST), IEEE, 2016, pp. 1–6.

26. A. Manichaikul, J. C. Mychaleckyj, S. S. Rich, K. Daly, M. Sale, W.-M. Chen, Robust relationship inference in genome-wide association studies, Bioinformatics 26 (22) (2010) 2867–2873.

27. D. O. Ricke, DNA mixtures from one or more sources and methods of building individual profiles therefrom, International Patent Application No. PCT/US2018/041081.

28. M. Conomos, A. Reiner, B. Weir, T. Thornton, Model-free estimation of recent genetic relatedness, Am J Hum Genet 98 (1) (2016) 127–148.

29. K.-H. Cheung, M. V. Osier, J. R. Kidd, A. J. Pakstis, P. L. Miller, K. K. Kidd, ALFRED: an allele frequency database for diverse populations and DNA polymorphisms, Nucleic Acids Res 28 (1) (2000) 361–363.

30. Mutation rates in humans. II. sporadic mutation-specific rates and rate of detrimental human mutations inferred from hemophilia B, Am J Hum Genet 64 (6) (1999) 1580–1587.

31. D. O. Ricke, A. Shcherbina, A. Michaleas, P. Fremont-Smith, GrigoraSNPs: Optimized HTS DNA forensic SNP analysis, J Forensic Sci. In press.

32. Ø. Skare, N. Sheehan, T. Egeland, Identification of distant family relationships, Bioinformatics 25 (18) (2009) 2376–2382.

33. T. Thornton, H. Tang, T. Hoffmann, H. Ochs-Balcom, B. Caan, N. Risch, Estimating kinship in admixed populations, Am J Hum Genet 91 (1) (2012) 122–138.

